# Anaerobic microbiota facilitate pathogen access to the airway epithelium in a novel co-culture model of colonization

**DOI:** 10.1101/2025.06.18.660211

**Authors:** Patrick J. Moore, Leslie A. Kent, Ryan C. Hunter

## Abstract

Chronic rhinosinusitis (CRS) is a prevalent condition characterized by mucus stasis, persistent inflammation, and infection of the paranasal sinuses. It often involves the bacterium *Pseudomonas aeruginosa,* especially in individuals with cystic fibrosis or a history of antibiotic use. While *P. aeruginosa* is a known opportunistic pathogen that employs a diverse array of virulence factors to cause airway infections, its ability to thrive in the sinonasal environment is also likely influenced by its local microbial ecology. For instance, anaerobic bacterial genera, such as *Streptococcus, Veillonella,* and *Prevotella,* are also commonly found in CRS and may contribute to *P. aeruginosa* persistence. Here we sought to test the hypothesis that anaerobes promote *P. aeruginosa* colonization of the upper airways, specifically through degradation of mucin glycoproteins that decorate the epithelial surface. Using a novel dual oxic-anoxic culture platform termed DOAC, we co-cultured Calu-3 epithelial cells with a CRS-derived anaerobic microbial community. We observed increased expression of inflammatory marker genes and degradation of mucin glycoproteins, along with enhanced *P. aeruginosa* colonization of the epithelial surface after anaerobe pre-treatment. Furthermore, mucins isolated from anaerobe-treated Calu-3 cells promoted greater *P. aeruginosa* attachment to microtiter plates *in vitro* compared to intact mucins. These results suggest that anaerobic microbiota may shape the sinonasal environment in a way that favors *P. aeruginosa* persistence, offering new insights into CRS pathogenesis and potential therapeutic targets to disrupt detrimental bacterial interactions in chronic airway disease.

**IMPORTANCE:** The prevalence and abundance of strict and facultative anaerobic bacteria in chronic sinusitis as detected by culture-independent sequencing has renewed interest in their potential role(s) in disease onset, progression, and treatment. However, reductionist study of interactions between anaerobic microbiota and the host has been limited by the lack of laboratory models compatible with their conflicting oxygen demands. The significance of this work lies in the use of a novel co-culture platform, termed DOAC, to interrogate anaerobe interactions with the airway epithelium. We use this platform to show that anaerobic CRS microbiota elicit a pro-inflammatory response, degrade mucin glycoproteins, and promote mucosal colonization by the canonical CRS pathogen, *Pseudomonas aeruginosa*. This work highlights highlight a potential role of anaerobic microbiota in conditioning the sinonasal environment to favor pathogen colonization.

## INTRODUCTION

Chronic rhinosinusitis (CRS) is a widespread and debilitating condition marked by persistent infection and inflammation of the paranasal sinuses(*1–3*). Among bacteria linked to CRS pathology, *Pseudomonas aeruginosa* is the second most common isolated pathogen in CRS(*4*) and is frequently associated with severe and refractory cases(*5*), particularly in people with cystic fibrosis (pwCF) or prior antibiotic exposure(*6–10*). While *P. aeruginosa* is a well-established opportunistic pathogen, its ability to successfully colonize the sinonasal environment is likely shaped by complex interactions with co-colonizing microbiota.

Strict and facultative anaerobic bacterial genera (*Streptococcus, Veillonella, Prevotella, Fusobacterium, Porphyromonas)*, which are also both prevalent and abundant among sinonasal microbiota(*9–11*), have increasingly been recognized as potential facilitators of *P. aeruginosa* persistence and pathogenicity(*12–15*). Previous studies suggest that anaerobes and their metabolites enhance *P. aeruginosa* biofilm formation(*13, 16, 17*), expand nutrient accessibility(*12, 18, 19*), alter antimicrobial susceptibility of airway pathogens(*20–23*), and modulate immune responses in ways that may favor *P. aeruginosa* persistence(*24*). Although these findings have primarily been observed in lower airway diseases like cystic fibrosis, it remains unclear whether anaerobic bacteria promote *P. aeruginosa* colonization of the sinuses, and if so, what specific molecular mechanisms are involved.

A notable challenge in studying the anaerobe-pathogen-host dynamic is the lack of laboratory models that allow co-culture of oxygen-dependent epithelial cells and oxygen-sensitive anaerobes. To overcome this, we recently developed a dual oxic-anoxic culture platform (DOAC), in which polarized epithelial monolayers are cultured in an anoxic workstation while oxygen is supplied to the basolateral compartment (**Fig. S1**)(*25*). This setup creates a near-anoxic microenvironment on the apical surface, permitting co-culture of airway epithelial cells with strict and facultative anaerobes and the tractable evaluation of their interactions over time.

In this study, we use DOAC to evaluate the role of CRS-associated anaerobes (e.g., *Streptococcus, Veillonella, Prevotella* spp.) in promoting *P. aeruginosa* colonization of the airway epithelium. Specifically, we test the hypothesis that anaerobes condition a mucosal microenvironment conducive to *P. aeruginosa* colonization through the modification and degradation of mucin glycoproteins. Using a CRS-derived anaerobic microbial community to challenge mucus-overproducing Calu-3 epithelial cells(*26–28*), we observed limited cytotoxicity but enhanced inflammatory gene expression, modification of low molecular weight mucins, and enhanced pathogen colonization with variations depending on the specific anaerobe. Understanding the molecular and ecological interactions that enable *P. aeruginosa* to establish itself and thrive in the mucus-rich, oxygen-limited upper airways could provide valuable insights into CRS pathogenesis and reveal new therapeutic strategies to disrupt bacterial synergy.

## RESULTS

### Anaerobic airway microbiota promote an inflammatory host response

The lack of tractable epithelial cell culture systems compatible with hypoxic or anoxic growth has limited our understanding of host-anaerobe interactions. Prior work has shown that culture supernatants of anaerobic bacteria elicit pro-inflammatory cytokine expression *in vitro* through mixed-acid fermentation and production of SCFAs(*18, 29, 30*). However, it is not yet known how the host responds to the physical presence of anaerobes at the airway epithelial interface. To address this knowledge gap, we used the DOAC model to assess the response of immortalized Calu-3 airway epithelial cells to co-culture with CRS-associated anaerobic microbiota.

We first utilized a defined anaerobic bacterial community (ABC) isolated from the upper airways of an individual with chronic sinusitis. This representative community was derived through anaerobic enrichment culturing of surgically-collected sinus mucus and was selected for its dominant bacterial taxa (*Veillonella, Prevotella, Streptococcus*) which are associated with chronic sinus disease (**Fig. 1A**). These genera are also known to degrade mucin glycoproteins that decorate the surface of Calu-3 cells, support pathogen growth through cross-feeding, and enhance *P. aeruginosa* pathogenicity both *in vitro* and *in vivo*(*12, 14, 15, 31*). After 3h of equilibration in the DOAC system (i.e. oxygenated basolateral compartment, anoxic apical surface), Calu-3 cells were apically challenged with ∼8 x 10^6^ colony forming units (CFUs) of the anaerobic community and incubated for an additional 24h. Colonization was confirmed using scanning electron microscopy, which revealed bacterial cells at the epithelial interface (**Fig. 1B**). Importantly, anaerobes (4 x 10^6^ CFUs) were recovered after 24h by washing with PBS and plating on Brain Heart Infusion agar (BHI)(**Fig. 1C**). Washing with Triton X-100 resulted in a 0.7-log increase in recovery (1.6 x 10^7^ CFUs), indicating both anaerobe growth at the epithelial surface and either robust attachment or invasion of host cells (i.e., bacterial cells were not removed by PBS washing alone). Recovery of ∼7 x 10^5^ CFUs on a *Prevotella-*selective medium (Brucella Blood Agar, BBA) under anaerobic conditions confirmed that apical oxygen concentrations were sufficiently low to support strict anaerobe growth, consistent with our previous work showing anaerobe proliferation and mixed-acid fermentation under DOAC conditions(*25*). Despite this bacterial growth on the apical surface of the epithelial cells, cytotoxicity was not induced by anaerobe challenge after 24h (**Fig. 1D**).

**Fig. 1.**
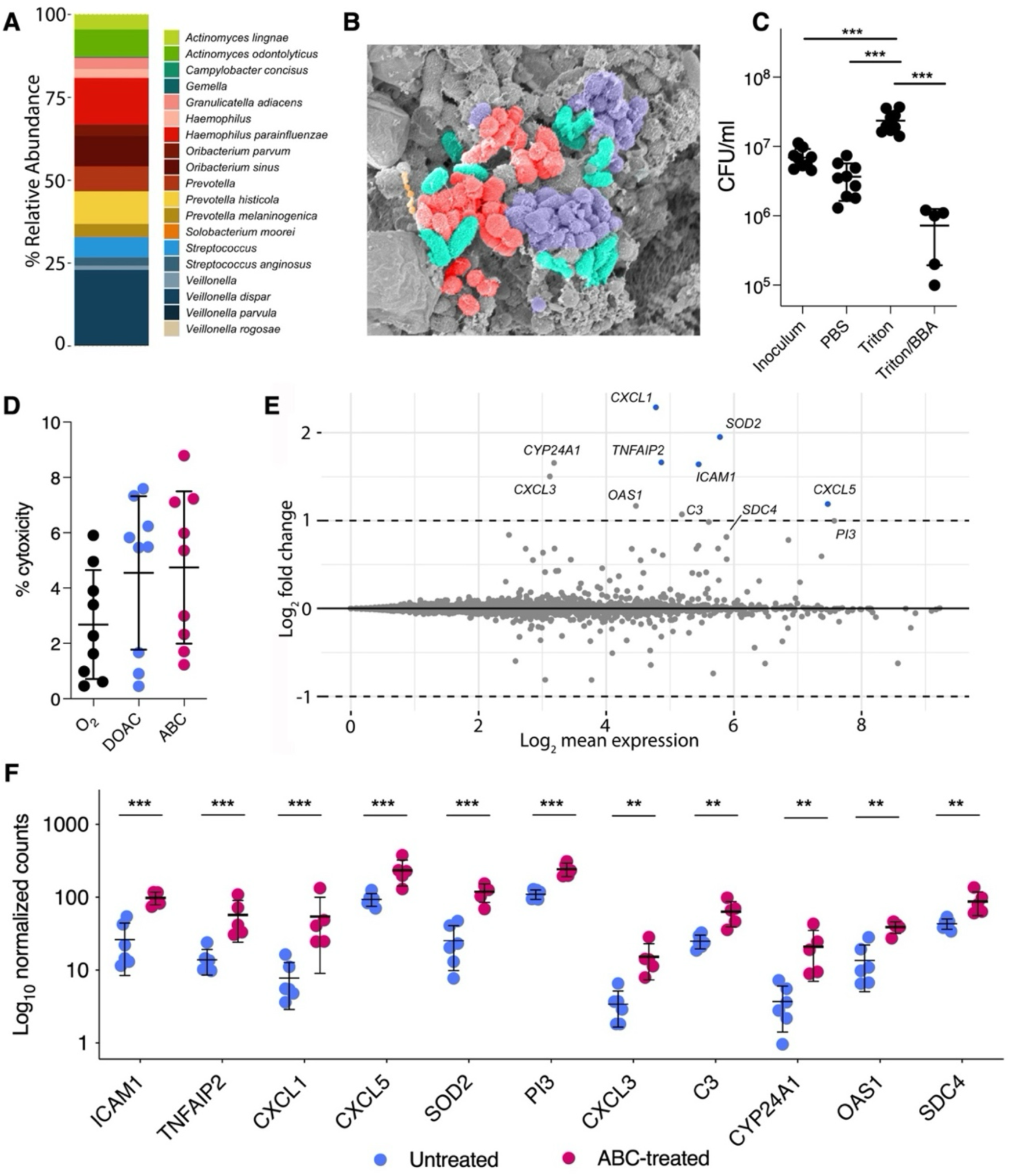
Anaerobic microbiota colonize the apical surface of Calu-3 cells and induce a pro-inflammatory response. **(A)** Taxonomic composition of an anaerobic bacterial consortium (ABC) derived from the upper airways. **(B)** SEM micrograph of Calu-3 cells after CRS challenge. **(C)** Bacterial recovery from Calu-3 cells after 24h by washing with PBS, TritonX-100, and plating on *Prevotella* selective agar (BBA). **(D)** Anaerobe (ABC) challenge did not result in Calu-3 cytotoxicity relative to unchallenged O2-grown or DOAC-grown cells. **(E)** MA plot representation of Calu-3 gene expression under ANLI after ABC challenge relative to an untreated ANLI control. **(F)** Log10-normalized gene counts from five or six independent biological replicates. Data in panels C and D were compared using a one-way ANOVA (*p*<0.0001) with multiple comparisons test. Data in panels E and F were compared using a Wald test, Benjamini-Hochberg adjusted (\*\*\**p*<0.001, **<0.01).

We then used RNAseq to profile the transcriptional response of Calu-3 cells to the anaerobe community (**Fig. 1E, Data S1**). Contrary to our expectations, only five genes showed differential expression relative to untreated cell cultures, all of which were upregulated (log2 fold change > 1, adjusted p-value <0.001). These genes were all markers of inflammation, and included *ICAM1*, *TNFAIP2* (which is regulated by TNFα in response to bacterial challenge)(*32*), chemokines *CXCL1* and *CXCL5* (which act as neutrophil chemoattractants and are primarily expressed during acute inflammation)(*33, 34*), and *SOD2* (superoxide dismutase 2, which is expressed in response to lipopolysaccharide and protects against apoptosis caused by inflammatory cytokines)(*35*). Additional inflammatory markers, such as *PI3* (elafin, an elastase inhibitor that primes innate immune responses in the lung)(*36*), *CXCL3*, *C3* (complement), *CYP24A1* (cytochrome p450 family 24 subfamily A member 1), *OAS1* (oligoadenylate synthetase), and *SDC4* (syndecan 4) were also differentially expressed, but did not reach statistical significance (**Fig. 1F**).

### Anaerobic microbiota alter the mucosal interface via mucin degradation

Previous work by us and others demonstrated the functional capacity of anaerobic microbiota to degrade airway mucins and support the growth of canonical pathogens via nutrient cross-feeding(*12, 31, 37*). Thus, in support of downstream pathogen colonization experiments, we used fast protein liquid chromatography (FPLC) to determine whether anaerobe challenge altered Calu-3 mucin integrity relative to unchallenged cells. To do so, we collected and purified mucin from the apical side of the Transwells as previously described(*38*) and used size-exclusion chromatography to assay their integrity. As expected, chromatograms revealed two characteristic peaks; (i) high molecular weight mucins which ran in the void volume of the column, and (ii) a broader inclusion volume peak representative of lower molecular weight mucins(*39, 40*) (**Fig. 2A**). While differences in the chromatographic profile of peak 1 (high molecular weight mucins) were negligible between culture conditions (*p*=0.38), peak 2 area was significantly reduced (*p*=0.007) following anaerobe challenge (**Fig. 2B**), reflecting degradation of lower-molecular weight mucin glycoproteins. These data suggest that in addition to eliciting an inflammatory host response, anaerobic colonization alters the physicochemical properties of mucin-decorated epithelial surface.

**Fig. 2.**
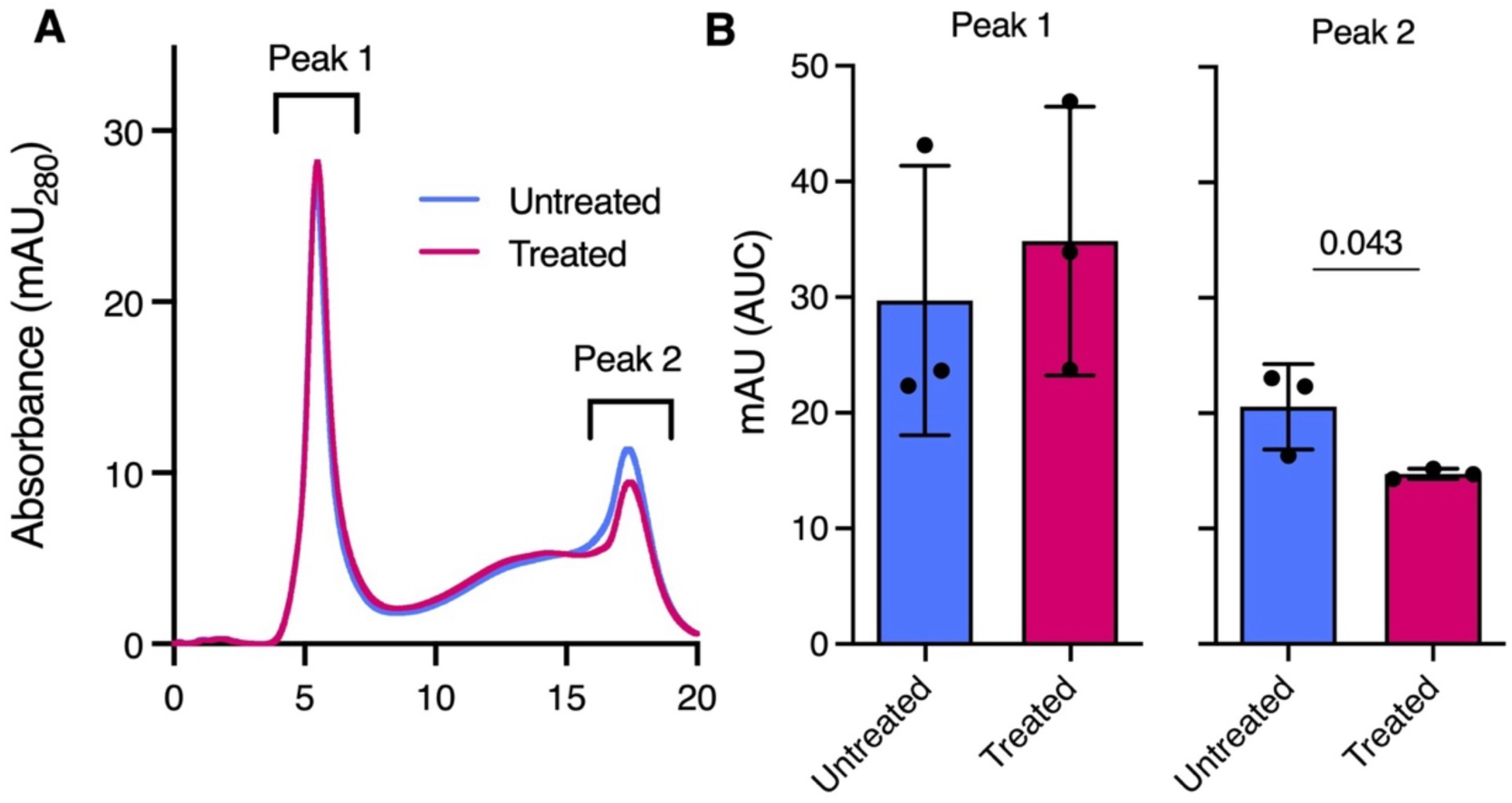
Anaerobic microbiota alter epithelial mucin integrity. **(A)** Representative FPLC traces of MUC5AC mucins purified from Calu-3 cells grown at ANLI (untreated) and after treatment with an anaerobic bacterial community (ABC, treated). **(B)** Area under curve (AUC) for both peak 1 (high molecular weight mucins) and peak 2 (low molecular weight mucin). Data shown were derived from three independent experiments using three biological replicates (n=9). Data were compared using a unpaired t-test with Welch’s correction (**, *P*<0.01).

### Anaerobes promote *P. aeruginosa* colonization of the airway epithelium

Recent studies have shown that viral challenge of the respiratory epithelium enhances *P. aeruginosa* colonization via interferon-mediated effects(*41*). Other work has demonstrated that *Streptococcus mitis* compromises the protective role of the mucus barrier by hydrolyzing mucin glycans(*42*). Given that the anaerobe consortium in our model both triggered an inflammatory response and altered mucin integrity, we hypothesized that, in addition to providing nutrients for pathogen growth through cross-feeding(*12*), anaerobic microbiota enhance *P. aeruginosa* colonization of the airway epithelium (illustrated in **Fig. 3A**).

**Fig. 3.**
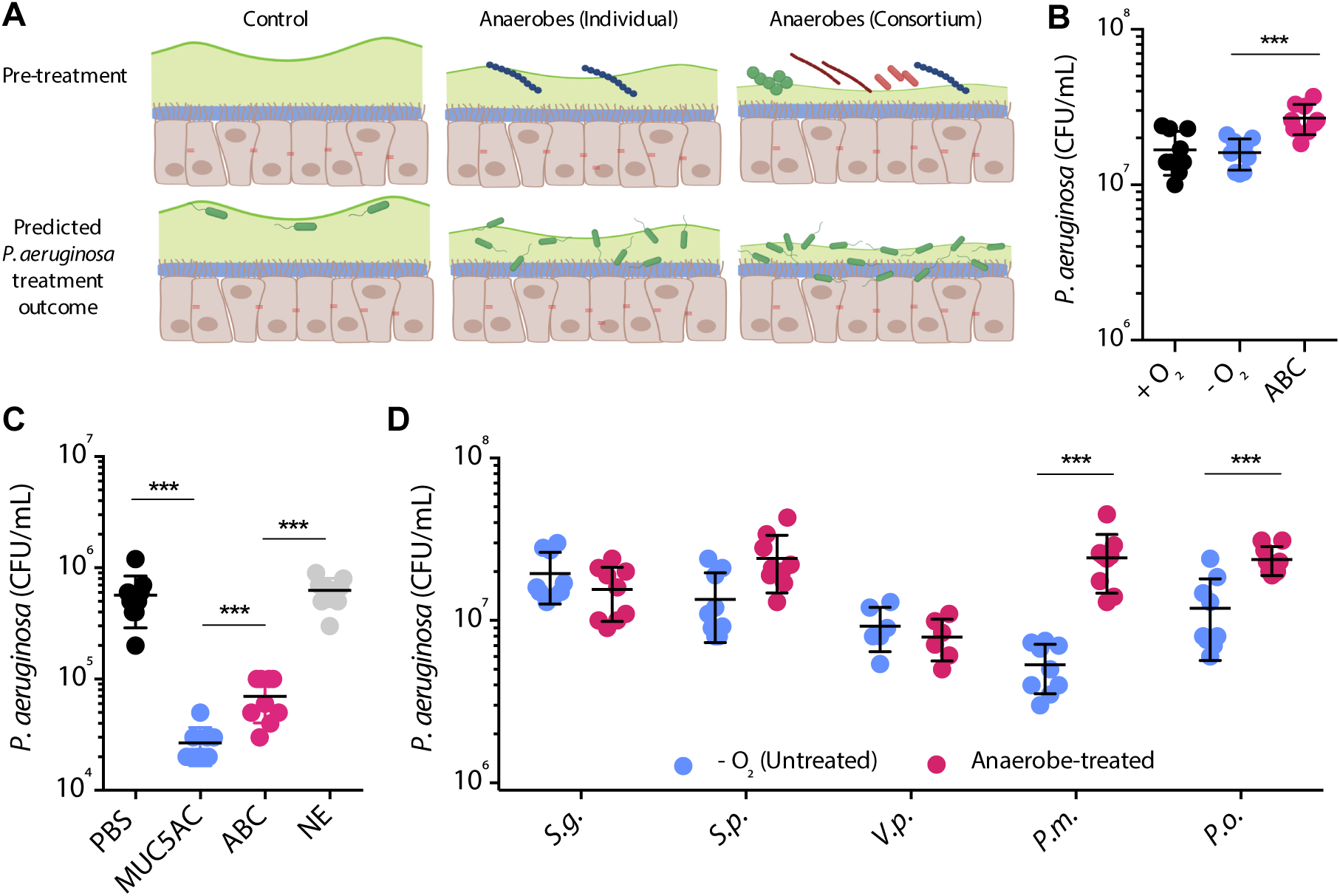
Anaerobic microbiota condition the epithelial interface for pathogen colonization. **(A**) Schematic of experimental design. **(B)** Calu-3 pre-treatment with an anaerobic bacterial consortium (ABC) potentiates *P. aeruginosa* colonization. **(C)** *P. aeruginosa* adhesion to microtiter plates coated with mucin (MUC5AC), ABC-treated mucin, and neutrophil elastase (NE)-treated mucin relative to an uncoated control (PBS). **(D)** *P. aeruginosa* adhesion to Calu-3 cells after pre-treatment with individual anaerobes (*S.g., Streptococcus gordonii; S.p., S. parasanguinis; V.p., Veillonella parvula; P.o., Prevotella oris; P.m., P. melaninogenica)*. Data shown in panels B-D were derived from three independent experiments with three biological replicates (n=9). Data in C were compared using a one-way ANOVA (*P*<0.0001) with multiple comparisons. Data in C were compared using a non-parametric Kruskal-Wallace test (*P*<0.0001) with multiple comparisons. Pairwise comparisons in panel D were performed using were compared using an unpaired t-test with Welch’s correction (*, *P*<0.05; ***, *P*<0001).

To test this hypothesis, Calu-3 cells were first treated with the anaerobic consortium (shown **Fig. 1A**) for 24h, then washed to remove spent medium and unbound cells. The cells were subsequently infected with ∼5 x 10^7^ CFUs of *P. aeruginosa* PA14 for 2h. Removing apical culture media and limiting the incubation time ensured that any difference in colonization between the anaerobe-treated cells and untreated (i.e., no anaerobe) control was due to cell attachment and not enhanced pathogen growth over time. *P. aeruginosa* colonization was determined by washing and permeabilizing the Calu-3 cells, followed by plate enumeration. As predicted, anaerobe pre-treatment resulted in an ∼67% increase in *P. aeruginosa* CFUs (1.1 x 10^7^) relative to untreated controls (*p*=0.0003, **Fig. 3B**).

To further confirm that enhanced pathogen colonization was due, at least in part, to mucin degradation, mucins isolated from Calu-3 cells treated with anaerobes (and untreated controls) were used to coat the surface of a microtiter plate, followed by the addition of *P. aeruginosa* PA14. Consistent with recent work showing that mucins disperse *P. aeruginosa* biofilm growth(*43*), mucin coating led to a 1.5 log-reduction in bacterial attachment compared to uncoated plates (PBS alone), as expected. In contrast, anaerobe degraded mucins showed a significant increase (*p*=0.0007) in *P. aeruginosa* binding to the microtiter plate compared to untreated mucins (**Fig. 3C**). To ensure that our mucin purification process (e.g., using guanidine hydrochloride) did not affect *P. aeruginosa* viability, plates were also coated with mucins degraded with human neutrophil elastase (NE) and isolated using the same procedure. NE-treated mucins led to similar PA14 attachment to PBS controls, confirming bacterial viability and showing that anaerobic microbiota can enhance pathogen colonization of a mucin-coated surface.

Finally, to assess the contributions of individual anaerobes to *P. aeruginosa* colonization, we challenged Calu-3 cells with representative isolates of the three most abundant genera in the anaerobic consortium (*Streptococcus, Veillonella, Prevotella)* prior to *P. aeruginosa* colonization (**Fig. 3D**). Contrary to a recent study(*42*), we found that *Streptococcus* species (*S. gordonii* and *S. parasanguinis*) had little effect on PA14 colonization, despite their known mucin degradation capacity(*44*). Similarly, *V. parvula* resulted in no significant differences between treatment conditions. In contrast, challenge with both *P. melaninogenica* and *P. oris,* two species commonly associated with inflammatory airway disease, led to significantly increased PA14 recovery from Calu-3 cells compared to unconditioned controls (*p=*0.0003 and *p=*0.0004, respectively). These data demonstrate that while anaerobic microbiota of the upper airways facilitate enhanced pathogen colonization of the epithelial surface, this effect is species-specific.

## DISCUSSION

Anaerobic bacteria have long been detected in the upper airways by culture (*45–47*), but the advent of culture-independent sequencing has renewed interest in their role(s) in chronic sinusitis. While in vitro data support potential pathogenic mechanisms, their relevance remains unclear due to a lack of suitable anaerobe-host interaction models compatible with the conflicting oxygen demands between cell types. To address this, we developed an *in vitro* platform for extended co-culture of polarized airway epithelial cells with anaerobic microbiota(*25*). Using this model, here we show that anaerobes commonly associated with the oral cavity - *Streptococcus, Prevotella,* and *Veillonella spp.* – may help establish a sinonasal inflammatory environment that promotes *P. aeruginosa* growth(*41, 48*). Additionally, we demonstrate that these anaerobes enhance pathogen colonization of the airway epithelial surface by degrading mucin glycans.

Previously, we reported that anaerobes stimulate pathogen growth through mucin-based cross-feeding(*12, 31*). Since *P. aeruginosa* cannot efficiently catabolize mucins alone(*37*), it relies on anaerobe-mediated breakdown and fermentation of mucin polypeptides and O-linked glycans for access to bioavailable substrates. Though not directly tested here, it is plausible that pre-colonization with anaerobes liberates mucin-derived metabolites, supporting pathogen persistence at the epithelial interface. While selective degradation of low molecular weight mucins remains unclear, our FPLC data confirm that anaerobic colonization structurally modifies the apical surface. Given that mucin degradation weakens barrier function and promotes pathogen-epithelial interactions(*42*), we propose that beyond the canonical mucus stagnation that accompanies CRS, anaerobic microbiota contribute to disease morbidity through a multifactorial process involving inflammation, cross-feeding, and mucosal surface alteration.

Focusing on an early time point post-*P. aeruginosa* challenge (2h), we evaluated anaerobic mucin degradation and its role in reshaping the epithelial interface to enhance pathogen attachment, as seen previously with *S. mitis* and *Neisseria meningitidis*(*42*). As expected, anaerobe degradation of apical mucus significantly increased *P. aeruginosa* attachment, which has important clinical implications. For example, epithelial colonization accelerates *P. aeruginosa* biofilm maturation, increases extracellular polysaccharide production, and induces quorum-sensing and other transcriptional changes(*49*). Additionally, *P. aeruginosa* grown on bronchial epithelial cells is far more resistant to antibiotics than when cultured on abiotic surfaces(*49*), consistent with its increased tolerance *in vivo*. We posit that anaerobe-mediated colonization further potentiates these phenotypes. Future studies will be needed to evaluate *P. aeruginosa-*host interactions over longer time periods to further understand differences in pathogen physiology and the inflammatory host response in the presence of anaerobic microbiota.

In conclusion, our findings highlight the potential role of anaerobic microbiota in shaping the sinonasal environment to favor *P. aeruginosa* colonization. By degrading mucin glycans and altering the epithelial interface, anaerobes may facilitate pathogen attachment and persistence, contributing to the onset and progression in chronic sinusitis and its recalcitrance to antibiotic therapy. Given the well-documented challenges of treating *P. aeruginosa* infections(*50*), particularly in biofilm-associated states, understanding the interplay between anaerobes, canonical airway pathogens, and the host will be critical. We are currently evaluating the mechanistic basis of anaerobe-mediated barrier disruption and potential therapeutic strategies to mitigate their impact on airway disease.

## METHODS

### Epithelial Cell Culture

Calu-3 cells were cultured in Minimal Essential Medium (MEM; Corning, USA) supplemented with 10% fetal bovine serum (FBS, Gene) and antibiotics (100 U/mL penicillin and 100 μg/mL streptomycin; Gibco) at 37°C in a 5% CO_2_ incubator. Once the cells reached 80% confluency, 1 x 10^5^ cells were seeded onto 6.5mm culture inserts (0.4 μm pore; Corning). After ∼5 days, apical medium was removed to establish an air-liquid interface (ALI). Cells were then maintained for an additional 21-28 days to allow differentiation and mucus accumulation.

To assemble the cell cultures in a gas-permeable multi-well plate manifold (DOAC)(Fig. S1), fully polarized Calu-3 cells were placed into a custom 3D-printed gasket fitted within a 24-well gas-permeable plate (CoyLabs, Grass Lake, MI) containing 800µL of MEM per well. Sterile mineral oil (400μL) was added to unused wells to prevent unwanted gas exchange. aThe assembled system was transferred into an anaerobic chamber (90% N_2_/ 5% H_2_/ 5% CO_2_), while a mixed gas flow (21% O_2_/ 5% CO_2_/ 74% N_2_) was supplied to the plate base to oxygenate the basolateral compartment. A schematic of this setup is provided in **Fig. S1**.

### Fast Protein Liquid Chromatography

Secreted mucins were collected from Calu-3 cells as previously described(*38*). Briefly, cells grown on Transwell inserts were solubilized in a reduction buffer containing 6M guanidine hydrochloride, 0.1M Tris-HCl, and 5mM EDTA (pH 8). To minimize mucin degradation, 10mM dithiothreitol (DTT) and a cOmplete Mini protease inhibitor tablet (Roche) were added to 400mL of reduction buffer before solubilization. Cell suspensions were gently agitated by pipetting to dislodge biomass, and suspensions from six Transwell inserts per plate were pooled into a single aliquot. Cells were rinsed with reduction buffer to remove residual mucin. This mixture was incubated at 37°C for 5h, followed by the addition of 25 mM iodoacetamide and overnight incubation at room temperature. Mucins were then dialyzed (1000 kDa MWCO) against 1L of 4M GuHCl buffer containing 2.25mM NaH_2_PO_4_-H_2_O and 76.8 mM Na_2_HPO_4_ for 36h, with buffer exchanges every 12h.

To evaluate the integrity of high-molecular-weight mucins, size-exclusion chromatography was performed using an Äkta Pure FPLC system (GE Healthcare BioSciences, Marlborough, MA) at 4°C. A 500 μL aliquot of purified mucin was manually injected and subjected to an isocratic run at a 0.4 mL/min flow rate for 1.5 column volumes using a 15mL 10/200 Tricorn column packed with Sepharose 4B-CL beads. The mobile phase consisted of 150mM NaCl in 50mM phosphate buffer (pH 7.2). Data were collected using Unicorn software (GE Healthcare Biosciences).

### Bacterial strains and culture conditions

*P. aeruginosa* PA14 was routinely cultured on Luria Bertani (LB) medium. The anaerobic strains *Prevotella melaninogenica* ATCC 25845, *Streptococcus parasanguinis* ATCC15912, and *Veillonella parvula* ATCC10790 were obtained from Microbiologics (St. Cloud, MN). *Streptococcus gordonii* was obtained from M.C. Herzberg (University of Minnesota) *and Prevotella oris* 12252T was purchased from the Japan Collection of Microorganisms. All anaerobes were maintained on Brain-Heart Infusion medium supplemented with 0.25 g/L hemin, 0.025 g/L vitamin K, and 5% (vol/vol) laked sheep’s blood (BHI-HKB) under anaerobic conditions. Additionally, a mucin-enriched anaerobic bacterial community (ABC) isolated from an individual with chronic rhinosinusitis and was enriched and cultured in a minimal mucin medium (MMM) as previously described(*12, 31*).

### RNA sequencing

The transcriptomic response of Calu-3 cells to anaerobe challenge was analyzed using RNAseq. Calu-3 cells were cultured at air-liquid interface and harvested after 24h of incubation under DOAC conditions with or without bacterial challenge. Normoxic (unchallenged) controls were maintained under standard incubator conditions. At the conclusion of each experiment, RNAlater (Invitrogen) was added to both the apical and basolateral compartments. RNA was extracted from 5 or 6 Transwells using the RNeasy Micro Plus kit (Qiagen) following the manufacturer’s instructions. DNase treatment was performed using the Zymo RNA Clean and Concentrator kit. RNA quality (RIN > 9.7) was assessed using an Agilent Bioanalyzer and RNA quantity was measured using RiboGreen. cDNA libraries were prepared using the SMARTer Universal Low Input RNA Kit (Takara Bio) and sequenced on an Illumina NovaSeq 6000 platform at the University of Minnesota Genomics Center.

For analysis, the Ensembl GTF annotation file was filtered to exclude non-protein-coding features. Fastq files were subsampled to a maximum of 100,000 reads per sample, and data quality was assessed with FastQC. Raw reads were aligned to the *Homo sapiens* reference genome (GRCh38) with Ensembl release 98 annotations. Gene counts were generated using the ‘featureCounts’ function of the RSubread package(*51*). Differential expression analysis was performed using DESeq2 (v.1.28.1), where size factors were estimated, gene-wise dispersions were computed, shrinkage was applied (type = ‘ashr’), and Wald hypothesis testing was conducted(*52, 53*). Genes with a log_2_ fold-change > 1 and a Benjamini-Hochberg adjusted *p*-value < 0.001 were considered significant. All code and data are available at (will be made available upon acceptance).

### Scanning electron microscopy

Untreated, anaerobe-challenged, and *P. aeruginosa*-infected cell cultures were washed three times in 0.2M sodium cacodylate buffer before fixation in a primary fixative solution (0.15 M sodium cacodylate buffer, pH 7.4, 2% paraformaldehyde, 2% glutaraldehyde, 4% sucrose, and 0.15% alcian blue 8GX) for 22h. Transwell membranes were washed three more times and treated with secondary fixative (1% osmium tetroxide, 1.5% potassium ferrocyanide, 0.135M sodium cacodylate, pH 7.4) for 90 minutes. After three additional washes, cells were dehydrated in a graded ethanol series (25%, 50%, 75%, 85%, 2 x 95%, and 2 x 100%) for 10 minutes per step, followed by CO_2_-based critical point drying using a Samdri-780A instrument (Tousimis, Rockville, MD). Transwell membranes were then mounted on SEM specimen stubs using carbon conductive adhesive tape and sputter coated with ∼5 nm iridium using the Leica ACE 600 magnetron-based system. Imaging was performed on a Hitachi S-4700 field emission SEM at an operating voltage of 2kV. Images were false colored using Adobe Photoshop CS6.

### Microtiter plate binding assay

The adhesion of *P. aeruginosa* adhesion to mucus was tested using an established microtiter plate-based assay(*54*). 96-well MaxiSorp microtiter plates (Nunc) were coated with 40μg/mL of mucins (MUC5AC) derived from untreated and ABC-treated Calu-3 cells. As a control, MUC5AC treated with neutrophil elastase (1 μg/mL) was also used. After coating, plates were incubated for 24h at 37°C. Mucin-coated wells were then washed three times with sterile PBS to remove any residual unbound mucin. Next, 5 x 10^7^ CFUs of *P. aeruginosa* PA14 were added to mucin-coated wells and incubated for an additional 2h at 37°C. Wells were washed 10 times with PBS to remove any unbound bacteria. Bound PA14 was desorbed by treating the wells with 200μL of 0.25% Triton X-100 for 15 minutes at room temperature. Bacteria bound to each well were enumerated by plating serial dilutions on LB agar. All assays were performed using three biological replicates and three technical replicates.

## ACKNOWLEDGEMENTS

Microscopy was performed in the University of Minnesota Characterization Facility (CharFac), which receives partial support from the NSF through the MRSEC (MR-2011401) and NNCI (ECCS-2025124) programs. P.J.M. was supported by a Cystic Fibrosis Foundation postdoctoral fellowship (MOORE18F0), L.A.K. was supported by an American Society for Microbiology Undergraduate Research Fellowship. This work was also supported by the Gus and Marjorie Esselen Cystic Fibrosis Trust Fund, a Gilead Sciences Investigator Sponsored Award to R.C.H., the National Heart Lung Blood Institute (R01136919), and the National Institute for Allergy and Infectious Diseases (R01HLAI177613).

