## Supplemental Data for "Anaerobic microbiota facilitate pathogen access to the airway epithelium in a novel co-culture model of colonization"

**Document S1.** Figure S1

**Data S1.** Excel file containing RNAseq data from Calu-3 cells challenged with CRS anaerobes for 24h

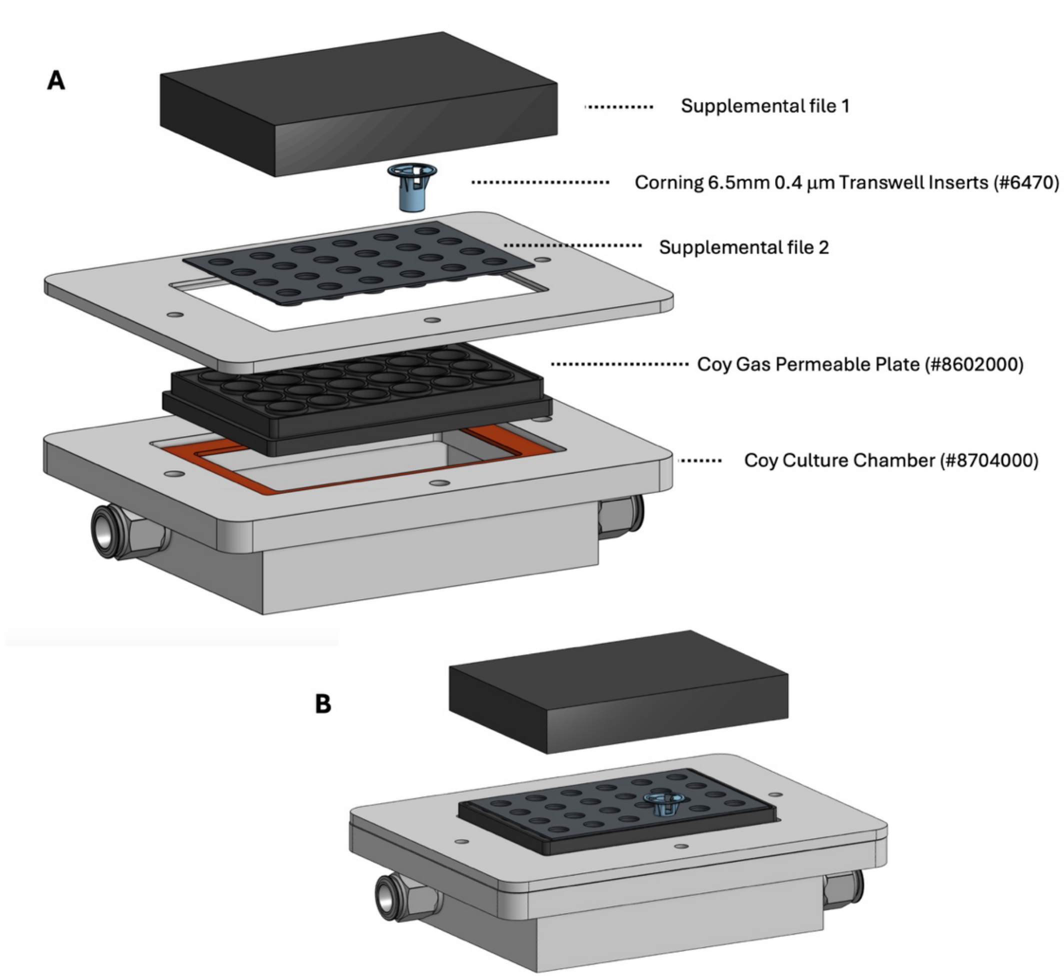


**Figure S1.** Schematic of an **(A)** unassembled and **(B)** assembled DOAC platform. Transwells are mounted in a custom 3D-printed thermoplastic polyurethane gasket mounted on a gas permeable multi-well plate. Blood gas is delivered and removed through cable glands mounted to the basolateral compartment of the Transwell-containing apparatus. (From Moore et al., 2025, mBio)
